# Physical Activity Delays Obesity-Associated Pancreatic Ductal Adenocarcinoma in Mice and Decreases Inflammation

**DOI:** 10.1101/2023.01.03.521203

**Authors:** Valentina Pita-Grisanti, Kelly Dubay, Ali Lahooti, Niharika Badi, Olivia Ueltschi, Kristyn Gumpper-Fedus, Hsiang-Yin Hsueh, Ila Lahooti, Myrriah Chavez-Tomar, Samantha Terhorst, Sue E. Knoblaugh, Lei Cao, Wei Huang, Christopher C. Coss, Thomas A. Mace, Fouad Choueiry, Alice Hinton, Jennifer M Mitchell, Rosemarie Schmandt, Michaela Onstad Grinsfelder, Karen Basen-Engquist, Zobeida Cruz-Monserrate

**Author notes:** Authors contributed equally (Valentina Pita-Grisanti, Kelly Dubay). **Correspondence:** Zobeida Cruz-Monserrate, Ph.D., Associate Professor, Department of Internal Medicine, Division of Gastroenterology, Hepatology and Nutrition, The Ohio State University Wexner Medical Center, 2041 Wiseman Hall, 400 W 12th Ave Columbus, OH 43210.

## Abstract

**BACKGROUND & AIMS:** Obesity is a risk factor for pancreatic ductal adenocarcinoma (PDAC), a deadly disease with limited preventive strategies. Lifestyle interventions to decrease obesity might prevent obesity-associated PDAC. Here, we examined whether decreasing obesity by increased physical activity (PA) and/or dietary changes would decrease inflammation in humans and prevent PDAC in mice.

**METHODS:** Circulating inflammatory-associated cytokines of overweight and obese subjects before and after a PA intervention were compared. PDAC pre-clinical models were exposed to PA and/or dietary interventions after obesity-associated cancer initiation. Body composition, tumor progression, growth, fibrosis, inflammation, and transcriptomic changes in the adipose tissue were evaluated.

**RESULTS:** PA decreased the levels of systemic inflammatory cytokines in overweight and obese subjects. PDAC mice on a diet-induced obesity (DIO) and PA intervention, had delayed weight gain, decreased systemic inflammation, lower grade pancreatic intraepithelial neoplasia lesions, reduced PDAC incidence, and increased anti-inflammatory signals in the adipose tissue compared to controls. PA had additional cancer prevention benefits when combined with a non-obesogenic diet after DIO. However, weight loss through PA alone or combined with a dietary intervention did not prevent tumor growth in an orthotopic PDAC model. Adipose-specific targeting of interleukin (IL)-15, an anti-inflammatory cytokine induced by PA in the adipose tissue, slowed PDAC growth.

**CONCLUSIONS:** PA alone or combined with diet-induced weight loss delayed the progression of PDAC and reduced systemic and adipose inflammatory signals. Therefore, obesity management via dietary interventions and/or PA, or modulating weight loss related pathways could prevent obesity-associated PDAC in high-risk obese individuals.

**Graphical Abstract:** 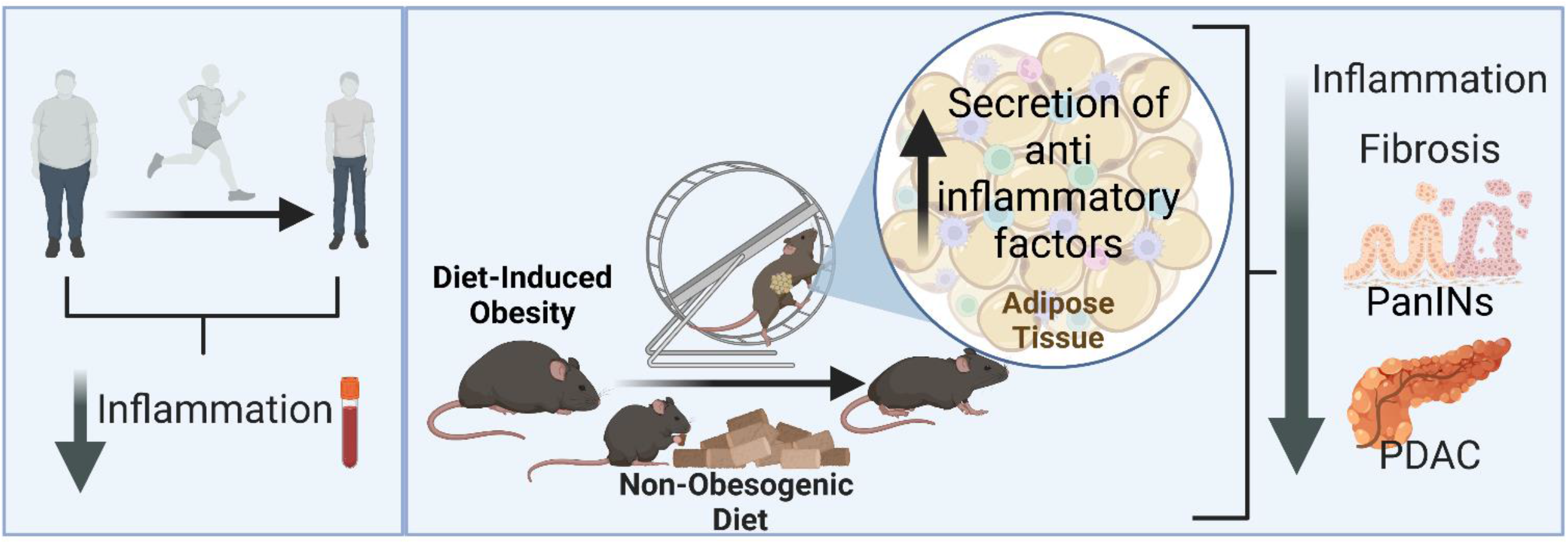

## INTRODUCTION

Pancreatic ductal adenocarcinoma (PDAC) is a deadly disease, with a 5-year survival rate of 11%^1^, mostly due to the lack of early detection, prevention, and treatment approaches^2^. Consequently, PDAC patients are mainly diagnosed at advanced stages of disease when surgical resection is often not possible^1, 2^.

Epidemiological studies indicate that premorbid obesity adversely influences PDAC mortality^3^ and increases the risk of PDAC^4–7^. High body mass index (BMI) is associated with early age of onset and decreased overall survival of PDAC^8^. This is concerning since 30% of the world’s population is overweight or obese and obesity rates are increasing due to western diets^9^. Obesity is associated with increases in systemic levels of inflammatory mediators, like interleukin 1 beta (IL-1β), interleukin-6 (IL-6), tumor necrosis factor-alpha (TNFα), and lipocalin-2 (LCN2), which can promote tumor growth^10–13^. Since obesity is a modifiable risk factor for PDAC^14, 15^, identifying effective lifestyle changes that reduce obesity or obesity-related inflammatory pathways could help minimize the incidence of obesity-associated PDAC^16^.

Lifestyle interventions that reduce obesity through diet alterations and increases in energy expenditure improve the outcomes of several diseases and overall health^17^. Studies have shown that moderate-to-vigorous physical activity (PA) reduces the risk of some cancers and chronic diseases^18–22^. PA also affects obesity by decreasing systemic inflammation while stimulating the release of beneficial cytokines, myokines, and/or adipokines^23^. PA can also reduce PDAC risk, with a stronger effect among younger populations^24^. Despite this, most PDAC clinical studies involving PA focus on increasing treatment efficacy and quality of life and do not evaluate the effects of increased PA or diet-related weight loss as a preventive strategy for obesity-associated PDAC^21^. We demonstrated that the diet-induced obesity (DIO) in mice accelerated pancreatic inflammation, fibrosis, pancreatic intraepithelial neoplasia (PanIN) lesions, and PDAC in a Kras-dependent manner^25^. Therefore, we hypothesize that implementing lifestyle interventions to decrease obesity through PA and/or diet modulation could prevent PDAC.

This study investigates the systemic levels of cytokines before and after a PA intervention in overweight and obese subjects and compares them to the changes observed in mice using similar interventions. We investigate whether increased PA, diet-induced weight loss, or both, alone or combined with chemotherapy, could prevent tumor progression, tumor growth, and inflammation in pre-clinical models of obesity-associated PDAC. We also explore whether targeting the expression of PA-associated anti-inflammatory signaling in the adipose tissue of mice delays tumor growth.

## MATERIALS AND METHODS

### Ethics Statement

All animal studies adhere to the Animal Research: Reporting of In Vivo Experiments (ARRIVE) guidelines^26^ and were compliant with the regulations and ethical guidelines for experimental animal studies of the Institutional Animal Care and Use Committee at The Ohio State University and the UTMD Anderson Cancer Center. The study on human subjects was approved by the Institutional Review Board of UTMD Anderson Cancer Center.

### PDAC Genetically Engineered and Orthotopic Mouse Models

All mice on PA were housed individually. *LSL-Kras^G12D^* mice^27^ were bred with the Ela-CreErT mice^28^ to generate inducible *Kras^G12D^*/CRE double transgenic mice (Kras^*G12D*^) and littermates (control) and given tamoxifen orally for 3 days as previously described^25^. Body weight (BW) was measured weekly. Blood collection was performed via submandibular bleeding and glucose was measured with Contour Next glucometer 7308 (Ascensia Diabetes Care, Parsippany, NJ).

Pancreatic tumor cells derived from a LSL-Kras^*G12D*^, LSL-Trp53^-/-^, Pdx1-CRE (KPC) genetically engineered mouse model (GEMM) transfected with enhanced firefly luciferase (KPC-LUC) were utilized in the PDAC orthotopic mouse model^12, 29, 30^. Cells were assessed for viability, counted, and mixed in PBS and 20% Matrigel (BD Biosciences, San Jose, CA). C57BL/6J mice (The Jackson Laboratories, Bar Harbor, ME) were orthotopically injected with KPC-LUC cells (0.25×10^6^ cells/mouse) as previously described^12, 30^, and given buprenorphine (0.05mg/kg subcutaneously every 12 hours for 48 hours, (Covetrus, Columbus OH). Subcutaneous injections of D-Luciferin (1.5mg/mouse; Caliper Life Sciences, Waltham, MA) were given to visualized tumor growth weekly using the In Vivo Imaging System (IVIS) (Caliper Life Sciences)^12, 30^. Bioluminescence was measured with the Living Image^®^ software. Body composition (BW and fat mass) was measured with the EchoMRI^™^ system (echoMRI LLC, Houston, TX).

### Voluntary Running Wheels

PA mice were provided access to a low-profile wireless running wheel while the no-PA control groups had access to a locked low-profile running wheel for mice (4 hours/day, 5 days/week) (Med Associates Inc, Fairfax, VT). The mice were placed in an altered light cycle environment (12:01 pm-12:00 am, noon-midnight dark cycle), for which they adjusted a week before starting interventions. Distance ran was tracked via a wireless USB hub (Med Associates Inc) with the Wheel Manager 2.03.00 acquisition software (Med Associates Inc).

### IL-15 Adipose-Specific AAV Targeting

An adipose-specific recombinant adeno-associated vector (rAAV) was generated containing dual expression cassettes that restrict off-target transduction in liver^31^.The first cassette consists of the CMV enhancer and chicken β-actin (CBA) promoter, woodchuck post-transcriptional regulatory element (WPRE) and bovine growth hormone poly-A flanked by AAV2 inverted terminal repeats as previously described^32^. The second cassette encodes a microRNA driven by the albumin basic promoter to target WPRE ^31^. Mouse IL-15 cDNA (GenBank: NM_001254747.4) was subcloned into a multiple cloning site downstream of CBA promoter as previously described^33^. The sequence was confirmed, and an empty vector was used as control. rAAV/empty and rAAV/IL-15 vectors were packaged into a Rec2 capsid. AAV viruses were purified by iodixanol gradient centrifugation as previously described^31, 34^ Female C57BL/6J mice received rAAV/empty or rAAV/IL-15 vectors (2×10^10^ vg per mouse via intraperitoneal injections of 100 μL in AAV dilution buffer). After 3 weeks of AAV injection, KPC-LUC cells were orthotopically implanted.

### Diet-Induced Obesity

Obesity (DIO) was induced using a high-fat diet (HFD), where 60% of the energy was derived from fat (DIO 58Y1 van Heek Series; Test Diet, St. Louis, MO) as previously shown^12, 25^. A non-obesogenic, lower-fat control diet (CD), where around 10% of the energy was derived from fat (DIO 58Y2; Test Diet), or (Pico Lab Rodent Diet 20 (5053); Lab Diet, St. Louis, MO) was used^25^.

### Histopathology and Pancreas Scoring

H&E sections were evaluated and scored by a board-certified veterinary pathologist (S.E.K.) blinded to genotype and study group. PanIN lesions were classified and graded according to standard criteria^35^. Pancreas were also scored for the most severe and frequent lesions in each section using a scoring scheme adapted from an existing grading scheme used for lesions in transgenic adenocarcinoma of the mouse prostate (TRAMP) mice ^36^. (**Supplementary Table 1**) Fibrosis and inflammation were assessed on a scale of 0 to 4. (**Supplementary Table 2**). The adjusted lesion scores and the fibrosis and inflammation scores generate the Total Pathological Score.

### Flow Cytometry Analyses of Immune Cell Populations

Mouse splenocytes were isolated and cryopreserved. Thawed splenocytes were re-suspended in 5% Fetal Bovine Serum PBS and cells (5× 10^5^/mL-1× 10^6^/mL) washed and incubated at 4°C with the fluorochrome-labeled antibodies and appropriate isotype controls. (**Supplementary Table 3**) Cells were washed, fixed in 1% formalin, and ran on a Fortessa (BD Biosciences) or Attune (Life Technologies, Carlsbad, CA) flow cytometers and analyzed with FlowJo^™^ software (BD Biosciences).

### Power In Motion Study

Obese/overweight subjects were recruited in the spring and fall of 2014 for a pilot study in partnership with the Power in Motion couch-to-5-kilometer training program (Houston, TX). Subjects participated in a 10-week training that included weekly talks and small group runs individually adapted. Blood samples and body composition using anthropometric measures and dual-energy x-ray absorptiometry (DXA) were compared before and after the program.

### Statistics

Statistical analyses were performed using the Prism 9 program (GraphPad Software San Diego, CA). A paired t-test was used for data of the same subjects before and after an intervention or paired Wilcoxon tests if non-normally distributed. Unpaired t-tests were used for comparing one variable between two groups or Mann-Whitney tests if non-normally distributed. Repeated measures two-way analysis of variance (ANOVA) with multiple comparisons using Sidak’s correction was used to compare two or more groups over time, or more than two variables. Log-transformation was performed for non-normally distributed data. One-way ANOVA test was used for comparing more than two groups, with Dunnett’s correction, or Kruskal Wallis test with Dunn’s correction for non-normally distributed data. Data are expressed as the mean ± standard error of the mean (SEM). Significance is displayed as \**P*≤0.05, \*\**P*≤0.01, \*\*\**P*≤0.001, \*\*\*\**P*≤0.0001.

## RESULTS

### PA decreases the expression of pro-inflammatory cytokines in overweight and obese human subjects

We assessed whether a PA intervention in overweight/obese subjects could reduce inflammation by analyzing serum levels of pro-inflammatory cytokines of unfit individuals who were overweight or obese before and after undergoing a 10-week running training program. After the intervention, subjects exhibited significantly decreased serum levels of the pro-inflammatory cytokines IL-8 (*P* = 0.002), LCN2 (*P* = 0.021), TNFα (*P* = 0.012), vascular endothelial growth factor (VEGF) (*P* = 0.027), interferon gamma (IFNγ) (*P* = 0.001), IL-17 (*P* = 0.005), macrophage inflammatory protein 1 alpha (MIP-1α) (*P* = 0.002), among others, compared to before the intervention (**Figure 1**). Moreover, the waist-to-hip ratio (WHR, *P* = 0.032) and waist circumference (*P* = 0.005) were significantly reduced, and there was a decreasing trend in % body fat (*P* = 0.060) but not BMI (*P* = 0.186) after the intervention (**Table 1**). These results confirm that a PA intervention can decrease systemic inflammation and modulate body composition in these subjects.

**Figure 1.**
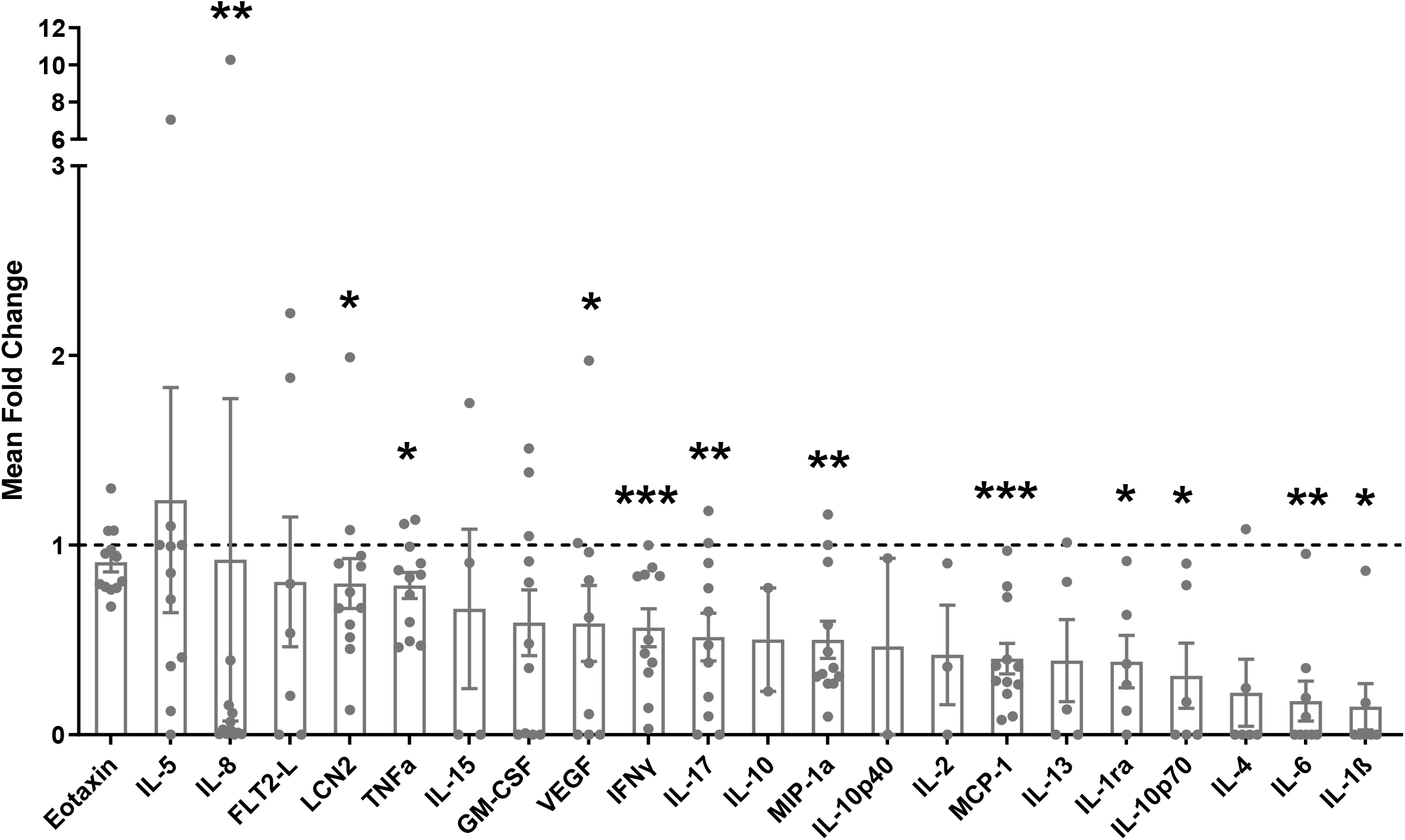
PA decreases the expression of pro-inflammatory cytokines in the serum of overweight/obese subjects. Mean fold change of serum cytokines after the program to before the program. Analyzed with a paired Wilcoxon test.

**Table 1.** Anthropometric measurements from overweight and obese subjects before and after a 10-week training program. Analyzed via paired t-test. **Abbreviations:** BMI = Body Mass Index, WHR = waist to hip ratio, FMI = Fat Mass Index, VAT = visceral adipose tissue.

|  | **Before**<br>**(n)** | **After**<br>**(n)** | **Before**<br>**Mean** | **After**<br>**Mean** | **Mean**<br>**Difference** | **Standard**<br>**Deviation of**<br>**Difference** | **T** | **P-**<br>**value** |
| --- | --- | --- | --- | --- | --- | --- | --- | --- |
| **Weight (kg)** | 18 | 12 | 80.73 | 76.71 | -0.53 | 1.07 | -1.73 | 0.112 |
| **BMI (kg/m2)** | 18 | 12 | 31.37 | 29.38 | -0.16 | 0.39 | -1.41 | 0.187 |
| **Waist (cm)** | 18 | 12 | 90.28 | 85.45 | -1.78 | 1.77 | -3.5 | **\*0.005** |
| **WHR** | 18 | 12 | 0.80 | 0.78 | -0.02 | 0.02 | -2.45 | **\*0.032** |
| **% Body Fat** | 18 | 12 | 43.41 | 42.29 | 0.35 | 0.58 | 2.09 | 0.060 |
| **FMI** | 18 | 12 | 13.91 | 12.65 | 0.04 | 0.24 | 0.54 | 0.598 |
| **VAT Area (cm2)** | 18 | 12 | 98.96 | 83.66 | -5.32 | 21.3 | -0.86 | 0.406 |

### PA delays weight gain, inflammation, fibrosis, PanIN progression, and PDAC incidence in an obesity-associated PDAC GEMM

Since PA decreased inflammation in overweight and obese human subjects, we evaluated whether PA could improve inflammation and delay PDAC in mice using a DIO model. Kras^*G12D*^ and littermate control mice were placed on a DIO intervention using a HFD and were divided into a PA (HFD + PA) or DIO control intervention group (HFD) for 1 month (**Figure 2A**). The HFD + PA mice ran an average of 5 km/day, increasing the distance ran each week (*P* < 0.0001) (**Figure 2B**). Kras^*G12D*^ mice in the HFD + PA group tended to run more per day than the HFD control mice (*P* = 0.072) (**Figure 2B**). During the PA intervention, the HFD + PA mice from the control and Kras^*G12D*^ groups displayed attenuated weight gain compared to the HFD only intervention (**Figure 2C**). At the end of the study, mice that underwent the PA intervention gained less weight (Kras^*G12D*^ 4.62 ± 0.69 g, control 5.25 ± 0.93 g) compared to the HFD controls (Kras^*G12D*^ 6.38 ± 1.58 g, control 7.96 ± 1.5 g), but these differences were not statistically significant (**Figure 2D**). The pancreas weight and the non-fasting glucose levels remained similar between the groups (**Figure 2E and Supplementary Figure 1A**).

**Figure 2.**
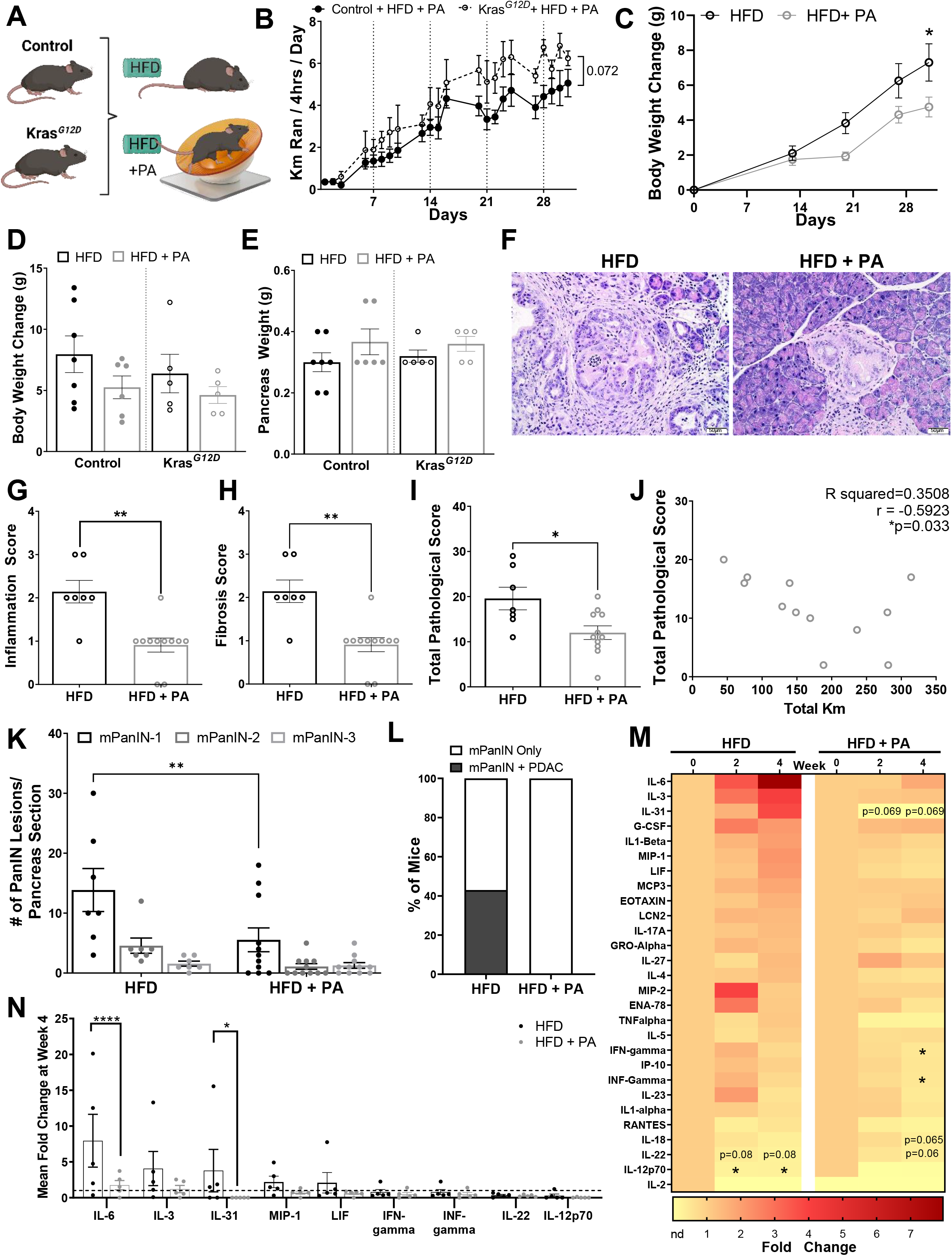
PA delays weight gain, inflammation, fibrosis, PanIN progression, and PDAC incidence in an obesity-associated GEMM of PDAC. (**A**) 40-day old Kras^*G12D*^ and control mice induced with tamoxifen on either a DIO intervention with HFD or a DIO intervention with HFD + PA for 31 days (n=7 control + HFD, n=5 Kras^*G12D*^ + HFD, n=6 control + HFD + PA, n=5 Kras^*G12D*^ + HFD+PA). (**B**) Average distance ran (km/day), analyzed via two-way ANOVA. (**C**) Average BW change over the 31 days, analyzed via two-way ANOVA. Significance displayed for the last day. (**D**) Average BW change and (**E**) pancreas weight after the intervention, analyzed with two-way ANOVA. (**F**) Representative H&E-stained pancreas sections of Kras^*G12D*^ mice (original magnification: x20, scale bar =50μm). Pathology scores for (**G**) inflammation, (**H**) fibrosis and (**I**) total pathology scores, based on the average of all H&E slides, analyzed with Mann-Whitney test. (**J**) Correlation between total pathological score and kilometers for the PA group and Pearson correlation coefficient. (**K**) Average number of PanIN lesions analyzed with two-way ANOVA (**L**) Percent of mice that developed PDAC. (**M**) Fold change serum cytokines of Kras^*G12D*^ mice at week 4 of intervention compared to baseline, analyzed with a paired Wilcoxon test. (**N**) Comparison of fold change serum cytokines at 4 weeks between HFD and HFD+PA groups, only displaying the cytokines with a trend or statistical significance in Figure 2M or had a *P* ≤ 0.999 at week 4, analyzed via two-way ANOVA.

The Kras^*G12D*^ mice in the HFD + PA group displayed significantly lower levels of pancreatic inflammation (*P* = 0.001), fibrosis (*P =* 0.001), and total pathological score (*P* = 0.027) compared to the HFD control group (**Figures 2F-I**). Additionally, there was a significant negative correlation between the total pathological score and the average distance ran (r = − 0.592; *P* = 0.033) indicating that increased running distance correlated with less advanced cancer (**Figure 2J**). Within the Kras^*G12D*^ mice, the HFD + PA group had significantly fewer PanIN-1 lesions (*P* = 0.002) than the HFD control group (**Figure 2K**). Notably, 43% of the mice in the HFD developed PDAC, while none of the mice in the HFD + PA developed PDAC (**Figure 2L**). Immunohistochemistry also showed that the pancreas of the HFD + PA mice had less fibrosis (αSMA), cell proliferation (Ki67), and macrophage infiltration (F4/80) compared to the HFD control group (**Supplementary Figure 1B**). We observed a trend for increased pro-inflammatory cytokines IL-6, IL-3, IL-31, granulocyte colony stimulating factor (G-CSF), IL-1β, MIP-1, leukemia inhibitory factor (LIF), MCP3, and LCN2 over time in the HFD control mice (**Figure 2M**). However, these trends were not statistically significant potentially due to large variability in the cytokine concentrations between mice and multiple comparisons. However, there were decreased levels of IL-12p70 (*P* = 0.019) and a decreasing trend in IL-22 (*P* = 0.08) (**Figure 2M**). Baseline levels of many pro-inflammatory cytokines were maintained in the PA intervention with significantly decreased IFN-gamma (*P* = 0.014) and INF-gamma (*P* = 0.014) and a trend towards decreased IL-31 (*P*=0.069); IL-18 (*P* = 0.065); and IL-22 (*P* = 0.06) (**Figure 2M**). When comparing the cytokines altered between the HFD and HFD + PA groups at week 4, most of the cytokines had a decreasing trend in the PA group, but IL-6 (*P* < 0.001) and IL-31 (*P* = 0.046) were significantly decreased (**Figure 2N**). These results demonstrate that PA delays the development of PDAC in a DIO GEMM, which might be mediated through anti-inflammatory effects, like how PA reduced inflammation in human subjects with obesity (**Figure 1**).

### Weight modulation by diet and/or PA delays PDAC development in an obesity-associated PDAC GEMM

We explored additional methods to decrease obesity by evaluating whether a combination of PA and diet modulation could further delay PDAC in DIO mice. *Kras^G12D^* and littermate control mice with DIO were assigned interventions for 7 weeks. One group continued on the DIO (HFD), the second group was switched to a non-obesogenic CD, and the third group was switched to a non-obesogenic CD with PA (CD + PA) (**Figure 3A**). All mice in the CD + PA group ran similar distances daily (~4.7 km/day) by the end of the intervention with an increase in distance ran each week (*P* < 0.001) (**Figure 3B**). Mice switched to the CD or the combined CD + PA interventions displayed significant weight loss during (**Figure 3C**) and at the end of the intervention (**Figure 3D**) (*P* < 0.05). Pancreas weight was significantly reduced in the Kras^*G12D*^ mice on the CD (0.33 ± 0.03g, *P* = 0.002) and the CD + PA (0.36 ± 0.03g, *P* = 0.013) interventions compared to the HFD control (0.59 ± 0.08g). In contrast, pancreas weight remained unchanged in the control mice after the interventions (**Figure 3E**). In the control mice both interventions (CD and CD + PA) decreased non-fasting glucose levels compared to the HFD group (*P* = 0.038 in CD, *P* = 0.003 in CD + PA) (**Figure 3F**). In the Kras^*G12D*^ mice non-fasting glucose levels were decreased in the CD + PA group compared to the HFD group (*P* = 0.035) (**Figure 3F**). These results suggest that reducing dietary fat intake drives weight loss, while PA synergistically enhances the effects of diet modulation in a DIO GEMM of PDAC.

**Figure 3.**
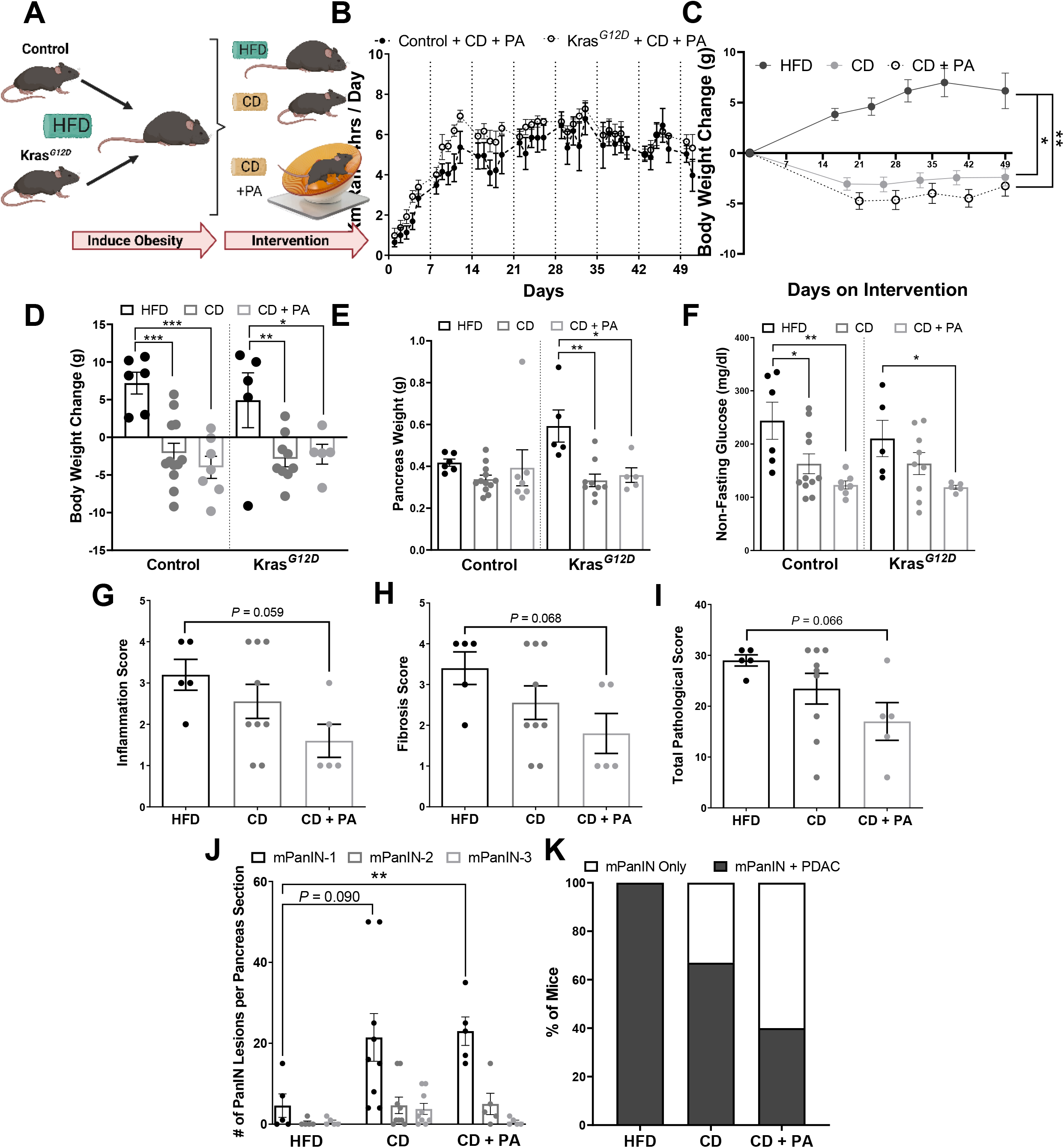
Diet-induced weight modulation and/or increased PA intervention reduces body weight, pancreas weight, non-fasting glucose levels and delays PDAC development in an obesity-induced GEMM of PDAC. (**A**) 40-day old *Kras^G12D^* mice were induced with tamoxifen for Kras activation and placed on a DIO intervention with a HFD for 33-54 days followed by 50 days with either no intervention (HFD) (control n = 6; Kras^*G12D*^ n = 5), CD (control n = 12; Kras^*G12D*^ n = 9), or CD+PA (control n = 7; Kras^*G12D*^ n = 5). (**B**) Distance ran by the PA mice, analyzed by a repeated measures two-way ANOVA. (**C**) BW change over time, analyzed via two-way ANOVA compared to HFD control. Significance displayed for the last day. (**D**) BW change from day 0 and (**E**) average pancreas weight, analyzed via two-way ANOVA. (**F**) Average non-fasting glucose at the end of the intervention, analyzed via two-way ANOVA after log transformation. Pathology score for (**G**) fibrosis and (**H**) inflammation and (**I**) Total pathology score, analyzed via Kruskal-Wallis test. (**J**) PanIN lesions analyzed via two-way ANOVA compared to the respective PanIN on the HFD control group. (**K**) Percent of Kras^*G12D*^ mice that developed PDAC at the end of each intervention.

Kras^*G12D*^ mice showed a decreasing trend of inflammation (*P* = 0.059), fibrosis (*P* = 0.068), and total pathological score (*P* =0.066) in the pancreas of the group that underwent the CD + PA intervention compared to the HFD group (**Figures 3G-I**). The mice in the HFD group had fewer PanIN-1 lesions than the CD (*P* = 0.090) or the combined CD + PA (*P* < 0.01) intervention groups (**Figure 3J**); however, 100% of the mice in the HFD group developed PDAC whereas only 67% in the CD and 40% in the CD + PA groups developed PDAC (**Figure 3K**). These findings illustrate the advantages of weight loss from a combination of dietary and PA interventions to delay tumor progression in an DIO **PDAC GEMM**.

### Diet and PA interventions reduce body weight but do not delay tumor growth in an obese orthotopic mouse model of PDAC

To determine whether diet and PA interventions could be preventive strategies to delay PDAC after obesity onset, C57BL/6J mice were exposed to DIO. Once obese, they were randomized to either a PA (HFD + PA), CD, or combined CD and PA (CD + PA) intervention or no intervention. After the interventions, KPC-LUC cells were implanted orthotopically while the mice continued on the dietary interventions without PA (**Figure 4A**). Mice in the HFD + PA and CD + PA interventions ran similar distances which increased over time (*P* < 0.0001) (**Figure 4B**). The HFD + PA had reduced weight (2.12 ± 0.836g, *P* = 0.002) and fat gain (0.745 ± 0.669g, *P* = 0.0003) before PDAC cell implantation compared to the mice in the HFD control group (6.88 ± 0.783g for weight; 5.609 ± 0.728g for fat) (**Figure 4C and Supplementary Figure 2A**). The mice in the CD and CD + PA interventions decreased weight (−4.4 ± 0.685g, *P* < 0.0001 for CD; −6.64 ± 0.796g, *P* < 0.001 for CD + PA) and body fat (−5.038 ± 0.751g, *P* < 0.0001 for CD; −7.977 ± 0.450g, *P* < 0.0001 for CD + PA) compared to the HFD control group before the PDAC cells were implanted (**Figure 4C and Supplementary Figure 2A**). These results suggest that a non-obesogenic diet intervention drives weight loss, but additional PA can still delay weight gain in the context of obesity.

**Figure 4.**
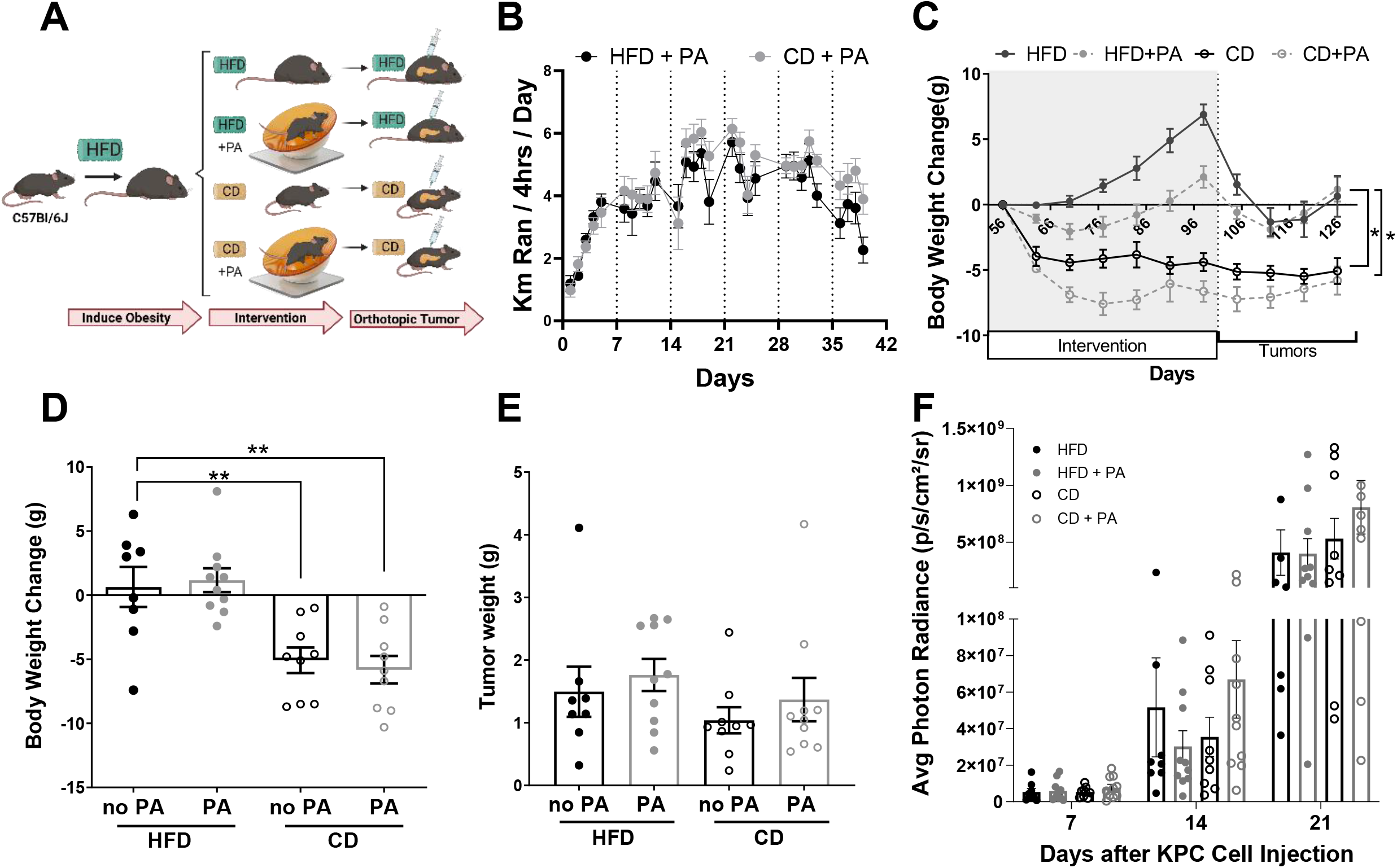
A diet and PA interventions reduce body weight but do not delay tumor growth in an obese orthotopic mouse model of PDAC. (**A**) 42-day old male C57BL/6J mice (n=10 per group) were placed on a DIO intervention with HFD for nine weeks before starting a 59-day CD and/or PA intervention. Mice were implanted with tumors and resumed their diet intervention. (**B**) Distance ran in kilometers analyzed via two-way ANOVA after log transformation. (**C**) BW change over time, analyzed with mixed-effects analysis comparing to HFD control. (**D**) Average change in BW from day 56 to the end of the intervention analyzed with two-way ANOVA. (**E**) Average tumor weight, analyzed via two-way ANOVA after log-transformation. (**F**) Average photon radiance signal from the tumor/week, analyzed after log-transformation via two-way ANOVA.

At study endpoint, BWs were significantly decreased in the CD (*P* = 0.0046) and CD + PA (*P* = 0.001) groups compared to the HFD control (**Figure 4D**), while the HFD + PA group had similar body weight changes to the HFD control. Tumor weights, and weekly tumor growth remained similar across all groups (**Figure 4E-F**). Interestingly, non-fasting glucose increased significantly in all the groups that underwent either a CD or PA, with the HFD group displaying the lowest levels of non-fasting glucose (**Supplementary Figure 2B**). These results show that neither PA nor a CD delayed tumor growth in a DIO orthotopic mouse model of PDAC.

We measured splenic immune cell populations to evaluate the effects of PA and diet on the immune system of the DIO PDAC orthotopic model. There were no significant differences between the percentages of MDSCs, F480^+^ cells, NK cells, CD8^+^ and CD4^+^ T cells between treatment groups (**Supplementary Figure 2C**), which aligned with the lack of changes in tumor growth. In contrast, when non-obese tumor-bearing mice were treated with gemcitabine in combination with PA (**Supplementary Figure 3A**), splenic CD8^+^ T cells were significantly elevated compared to a saline group without PA (*P* = 0.002) and a saline group with PA (*P* = 0.001) (**Supplementary Figure 3B**). We did not observe any changes in body weight (**Supplementary Figure 3C**) but did observe reduced body fat composition during the PA intervention (**Supplementary Figure 3D**). Although we saw a delay in early tumor growth due to PA compared to no PA in this model (*P* = 0.032) (**Supplementary Figure 3F**), we did not observe any additional benefits of a combined gemcitabine and PA intervention on tumor growth (**Supplementary Figure 3G**).

### PA promotes adipose-derived anti-inflammatory signaling that delays PDAC growth

To further understand how PA impacts inflammatory signaling in the adipose tissue, we performed a transcriptomic analysis of the adipose tissue from the Kras^*G12D*^ and control mice given a HFD or a HFD + PA intervention (**Figure 2A**) to assess changes in gene expression from the PA intervention (**Figure 5A**). PA increased the expression of genes involved in mRNA binding, amide and peptide biosynthesis, and mitochondria-related pathways, and decreased the expression of genes involved in phagocytosis and several immune-related pathways (**Figure 5A, 5B, and Supplementary Table 4**). Within these pathways, we observed a significant increase in the IL-15/IL-15 receptor agonist (ra) anti-inflammatory axis due to PA which we verified by quantitative RT-PCR (*P* = 0.003) (**Figure 5C**). Through viral targeting, we upregulated the expression of IL-15 in the adipose tissue of mice and then performed an orthotopic implantation of PDAC cells to measure tumor growth (**Figure 5D**). Viral targeted overexpression of IL-15 in the adipose tissue of the mice was verified by quantitative RT-PCR (*P* = 0.046) (**Figure 5E**). Tumor growth was significantly delayed in the mice with the adipose tissue-specific IL-15 upregulation compared to control mice (*P* = 0.0322) (**Figure 5F and G**). These results suggest that PA modulation of anti-inflammatory pathways in the adipose tissue significantly contributes to the benefits of PA as a prevention or treatment strategy for PDAC.

**Figure 5.**
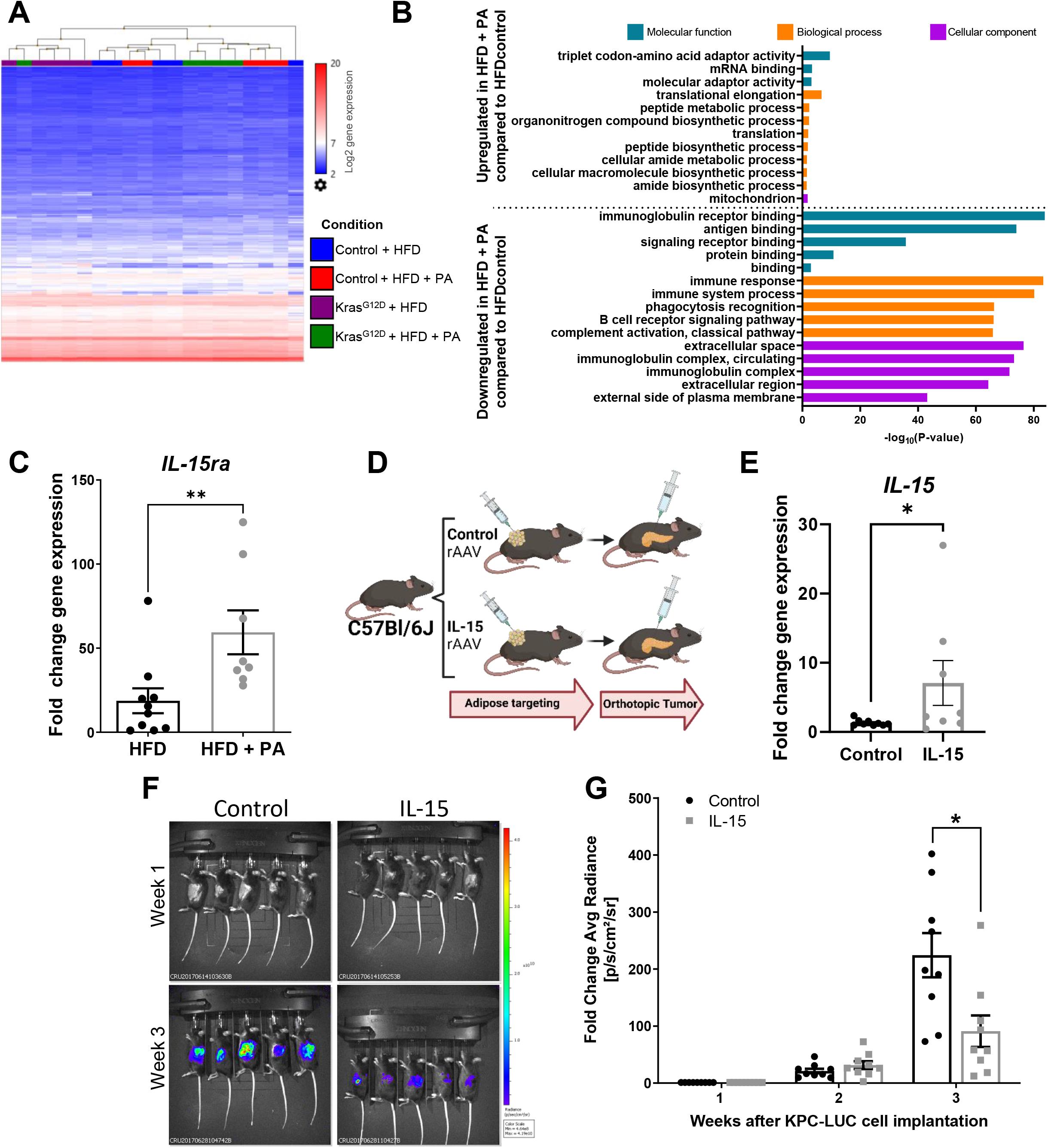
PA modulation of the anti-inflammatory pathways in the adipose tissue significantly delays PDAC tumor growth. (**A**) Heat map showing hierarchical clustering of genes from the adipose tissue of control and Kras^*G12D*^ mice after a DIO intervention of a HFD or HFD+PA from Figure 2A. Blue = low expression, red = high expression. (**B**) Gene ontology displaying top significant pathways modulated based on adipose tissue gene expression due to PA from Figure 2A cohort (**C**) Fold change adipose tissue gene expression of IL-15ra in control and Kras^*G12D*^ mice from Figure 2A, analyzed via Mann-Whitney test. (**D**) 9-week old Female C57BL/6J mice (n=10/group, 5/cage) received an empty/control or IL-15 adipocyte-targeting rAAV vector and were implanted with tumors 3 weeks later. (**E**) Fold change adipose tissue IL-15 gene expression at the end of the study, analyzed via Mann-Whitney test. (**F**) Representative IVIS images and (**G**) quantified fold change average photon radiance signal of mice with and without IL-15 adipose targeting, analyzed via two-way ANOVA after log transformation.

## DISCUSSION

Here, we report that a voluntary running-based PA intervention reduces systemic inflammation in overweight and obese subjects and in an obesity-associated PDAC GEMM. We also demonstrate that PA reduces pancreas fibrosis, macrophage infiltration, and delays PDAC in an obesity-associated PDAC GEMM. When PA is combined with a dietary intervention for weight loss, there is an additive beneficial effect to delay obesity-associated PDAC. However, when a HFD is given for a longer period prior to a PA and diet intervention in an orthotopic mouse model of PDAC, there are no changes in tumor growth. We also show that a PA intervention increases the expression of anti-inflammatory associated genes in the adipose tissue, like IL-15 and increasing IL-15 expression in the adipose tissue can be a strategy to delay PDAC tumor growth in mice. These findings support a prominent role for adipose-derived inflammation in the progression of obesity-associated PDAC and emphasize the potential of weight loss via PA and dietary changes to delay obesity-associated PDAC in obese high-risk individuals.

Epidemiological and animal studies show the benefits of weight loss interventions through PA in lowering the risk of several types of cancer and chronic diseases, including PDAC^19, 24, 38–40^. In PDAC patients, PA improved tumor vascularity and enhanced chemotherapy outcomes in a patient-derived xenograft mouse model^41^. Contrastingly, another study using a subcutaneous PDAC mouse model found that “low volume” continuous exercise reduced tumor growth, but this effect was not seen when the exercise was combined with chemotherapy^42^. We also demonstrated that PA had an initial delay in tumor growth but it did not enhance the effect of gemcitabine in a non-obese orthotopic mouse model of PDAC.

PA also improves glycemic control in PDAC patients after tumor resection^43^. Glucose control has become increasingly important for patients at a high risk of developing PDAC, since patients with new-onset diabetes have a ~8-fold higher risk of developing PDAC and can develop hyperglycemia before the tumor is identified^44, 45^. We demonstrated that the combination of a PA and diet intervention in an obesity-associated PDAC GEMM was successful at lowering blood glucose and could be a strategy to implement in at risk obese populations of PDAC to control glucose levels. However, in the obese orthotopic PDAC mouse model, glucose levels increased with the PA and diet interventions, which emphasizes the variations in outcomes based on the type of intervention and preclinical model utilized.

The impact of weight loss interventions on cancer outcomes might depend on the degree of chronic metabolic dysregulation that results from the amount of time consuming a HFD. Preclinical studies in rats found that long-term obesogenic diets increased hepatic oxidative stress and metabolic syndrome to a higher extent than short-term diets^46, 47^. In a mouse model of breast cancer, caloric restriction was more efficient at reverting the carcinogenic effects of obesity than a low fat diet consumption^48^. Here consumption of a HFD for longer than one month diminished the anti-tumorigenic effects of PA in PDAC and speculate that the extended time on a HFD before the CD or PA interventions might generate more permanent epigenetic or metabolic changes that hinders the benefits of PA seen in our one-month intervention. Further research is needed to investigate short- and long-term DIO with various modalities of weight loss interventions in PDAC.

Like our outcomes in the obese orthotopic PDAC model, weight normalization with a dietary intervention in an orthotopic breast cancer mouse model was insufficient to reverse obesity-associated inflammation and did not have any effect on tumor growth^10^. Likewise, a study in a patient-derived subcutaneous mouse model of PDAC did not find any differences in the tumor growth of mice after undergoing a treadmill running intervention post-tumor implantation^41^. Differences between the GEMMs, orthotopic and subcutaneous models (xenograft or allograft), as well as variations in the length, timing (pre- or post-tumor), and type of the intervention could contribute to the poor recapitulation between the different studies. The lack of tumor initiation steps in the orthotopic and subcutaneous mouse model eliminates any effect that the interventions have on the early stages of the disease. All of these variables need to be considered to answer prevention or treatment questions in cancer pre-clinically.

Other preclinical studies have found an immune cell-dependent effect of exercise in cancer^49^. Exercise increases CD8^+^ T cell infiltration and decreases the number of myeloid derived suppressor cells (MDSCs) suggesting that exercise can shift the tumor microenvironment from an immunosuppressive to an immunocompetent state, improving the effectiveness of immunotherapy^50–52^. Since PDAC is considered immunologically “cold” with low infiltration of CD8^+^ T cells, hindering cytotoxic effects in the tumor^53, 54^, increasing immune cell infiltration by PA represents a possible neoadjuvant for chemotherapies. Here, we did not see any PA-related changes in splenic immune cell populations in the context of obesity. We only saw an increase in splenic CD8^+^ T cells when PA was combined with gemcitabine treatments in non-obese mice. Since the PA intervention did not reduce tumor growth in our obese orthotopic mouse model of PDAC, it is possible that the PA intervention did not stimulate the immune cells to the extent needed to reduce tumor growth or that the effects of obesity are too strong to reverse in this model. In contrast, a recently published study that evaluated PDAC orthotopic tumors in combination with treadmill running, observed increased tumor infiltration of CD8^+^ T cells, decreased MDSCs, and a delay in tumor growth^55^. This same study found increased CD8^+^ T cells in the tumors of PDAC patients that underwent an exercise intervention before tumor resection^55^. In contrast, another study in a non-obese subcutaneous PDAC mouse model with various treadmill running regimes found that none of the interventions stimulated the immune system and that only low volume continuous exercise reduced tumor growth^42^. These running interventions were not in the context of obesity, which could explain the lack of tumor and immune effects seen in our model. Many PA preclinical studies in cancer differ in methodology utilized as we recently described^21^, which hinders the translation of the results to the clinic. Differences between forced treadmill running and voluntary wheel running are likely to have an impact on stress and other factors that influence the immune response and treatment outcomes^56^.

Apart from the effects of obesity, adipose tissue inflammation and secreted factors can worsen cancer outcomes^57^. For example, adipose tissue contributes to tumorigenesis by activating the CXCL12-CXCR4/CXCR7 signaling axis in prostate cancer^58^, increasing adipose fatty acid release in PDAC^59^, and increasing adipose-derived proinflammatory cytokines in breast cancer^60^. Activation of the IL15-IL15Rα-axis may also drive PA-induced immune anti-tumor effects^55^. Here, we show adipose-derived IL-15 is a prominent driver of anti-tumor effects, suggesting a major beneficial role for the adipose tissue molecular remodeling during PA. Nonetheless, more studies are needed to characterize the full extent of the adipose-tumor interactions and how weight loss interventions modulate it.

The findings of this study revealed that a PA and dietary intervention that lowers fat intake during cancer initiation and progression delay PDAC in DIO GEMMs, and these effects are likely mediated by a PA-induced anti-inflammatory effect in the adipose tissue. While additional studies are needed to better understand the mechanisms by which PA delays DIO PDAC progression, this study provides compelling evidence for a beneficial adipose-related antiinflammatory function of weight loss through PA and lower fat dietary interventions in obesity-associated PDAC.

## Abbreviations

CD: control diet
DIO: diet-induced obesity
ELISA: enzyme-linked immunosorbent assay
GEMM: genetically engineered mouse model
HFD: high fat diet
IL: interleukin
LCN2: lipocalin-2
MDSC: myeloid derived stem cells
NK: natural killer
PA: physical activity
PanIN: pancreatic intraepithelial neoplasia
PDAC: pancreatic ductal adenocarcinoma

## Conflict of interest/disclosures

None

## Author Contributions

1. Valentina Pita-Grisanti - study concept and design; development of methodology; acquisition of data; analysis and interpretation of data; drafting of initial manuscript; writing, review, and/or revision of the manuscript; administrative, technical, or material support; final approval of the version to be submitted
2. Kelly Dubay - study concept and design; development of methodology; acquisition of data; analysis and interpretation of data; drafting of initial manuscript; writing, review, and/or revision of the manuscript; administrative, technical, or material support; final approval of the version to be submitted
3. Ali Lahooti - study concept and design; development of methodology; acquisition of data; analysis and interpretation of data; writing, review, and/or revision of the manuscript; administrative, technical, or material support; final approval of the version to be submitted
4. Niharika Badi - development of methodology; acquisition of data; analysis and interpretation of data; writing, review, and/or revision of the manuscript; administrative, technical, or material support; final approval of the version to be submitted
5. Olivia Ueltschi - acquisition of data; analysis and interpretation of data; writing, review, and/or revision of the manuscript; administrative, technical, or material support; final approval of the version to be submitted
6. Kristyn Gumpper-Fedus - analysis and interpretation of data, drafting of initial manuscript, critical revision of the final manuscript, final approval of the version to be published
7. Hsiang-Yin Hsueh - analysis and interpretation of data, critical revision of the final manuscript, final approval of the version to be published
8. Ila Lahooti - acquisition of data; analysis and interpretation of data; writing, review, and/or revision of the manuscript; administrative, technical, or material support; final approval of the version to be submitted
9. Myrriah Chavez-Tomar - analysis and interpretation of data; drafting of initial manuscript; writing, review, and/or revision of the manuscript; administrative, technical, or material support; final approval of the version to be submitted
10. Samantha Terhorst - acquisition of data; writing, review, and/or revision of the manuscript; administrative, technical, or material support; final approval of the version to be submitted
11. Sue E. Knoblaugh - acquisition of data; analysis and interpretation of data; writing, review, and/or revision of the manuscript; administrative, technical, or material support; final approval of the version to be submitted
12. Lei Cao - acquisition of data; analysis and interpretation of data; writing, review, and/or revision of the manuscript; administrative, technical, or material support; final approval of the version to be submitted
13. Wei Huang - administrative, technical, or material support; final approval of the version to be submitted
14. Christopher C. Coss - writing, review, and/or revision of the manuscript; administrative, technical, or material support; final approval of the version to be submitted.
15. Thomas A. Mace - acquisition of data; analysis and interpretation of data; writing, review, and/or revision of the manuscript; administrative, technical, or material support; final approval of the version to be submitted.
16. Fouad Choueiry - acquisition of data; analysis and interpretation of data; writing, review, and/or revision of the manuscript; administrative, technical, or material support; final approval of the version to be submitted.
17. Alice Hinton - analysis and interpretation of data; writing, review, and/or revision of the manuscript; final approval of the version to be submitted.
18. Jennifer M Mitchell - acquisition of data; analysis and interpretation of data; writing, review, and/or revision of the manuscript; administrative, technical, or material support; final approval of the version to be submitted.
19. Rosemarie Schmandt - acquisition of data; analysis and interpretation of data; writing, review, and/or revision of the manuscript; administrative, technical, or material support; final approval of the version to be submitted
20. Michaela Onstad Grinsfelder - acquisition of data; analysis and interpretation of data; writing, review, and/or revision of the manuscript; administrative, technical, or material support; final approval of the version to be submitted
21. Karen Basen-Engquist - acquisition of data; analysis and interpretation of data; writing, review, and/or revision of the manuscript; administrative, technical, or material support; final approval of the version to be submitted
22. Zobeida Cruz-Monserrate - study concept and design; development of methodology; acquisition of data; analysis and interpretation of data; drafting of initial manuscript; writing, review, and/or revision of the manuscript; administrative, technical, or material support; final approval of the version to be submitted; study supervision.

## Data Transparency Statement

All data and analytic methods are included in the manuscript and supplementary material.

## Acknowledgements

We thank The OSUCCC Small Animal Imaging Core Laboratory for their In Vivo Imaging System and echoMRI services, Comparative Pathology and Digital Imaging Shared Resource for immunohistochemical support and histopathologic evaluation, Analytical Cytometry Shared Resource, and Genomics Shared Resource for their mRNA Affymetrix assay and sequencing support. Graphical abstract and experimental design figures were created with BioRender.com

## Notes

**Grant Support**: Research reported in this publication was supported by Knowledge GAP MDACC (ZC-M), Start-up funds from The Ohio State University Comprehensive Cancer Center (OSUCCC) (ZC-M). The intramural research program Pelotonia Idea Award from the OSUCCC (ZC-M), Pelotonia Scholarship Program (MC-T, KD, AL, OU, and VP-G). The National Center for Advancing Translational Sciences TL1TR002735 (KG-F), and in part by the MD Anderson Cancer Center Support Grant CA016672 and the OSUCCC P30 CA16058 National Cancer Institute. The content is solely the responsibility of the authors and does not necessarily represent the official views of the National Institutes of Health. Any opinions, findings, and conclusions expressed in this material are those of the author(s) and do not necessarily reflect those of the Pelotonia Scholarship Program, The Ohio State University, and the National Institutes of Health.

### Competing Interest Statement

The authors have declared no competing interest.

## References

1. Siegel RL, Miller KD, Fuchs HE, et al. Cancer statistics, 2022. CA Cancer J Clin 2022;72:7–33.

2. Zhao Z, Liu W. Pancreatic Cancer: A Review of Risk Factors, Diagnosis, and Treatment. Technol Cancer Res Treat 2020;19:1533033820962117.

3. Majumder K, Gupta A, Arora N, et al. Premorbid Obesity and Mortality in Patients With Pancreatic Cancer: A Systematic Review and Meta-analysis. Clin Gastroenterol Hepatol 2015.

4. Genkinger JM, Spiegelman D, Anderson KE, et al. A pooled analysis of 14 cohort studies of anthropometric factors and pancreatic cancer risk. Int J Cancer 2011;129:1708–17.

5. Kasenda B, Bass A, Koeberle D, et al. Survival in overweight patients with advanced pancreatic carcinoma: a multicentre cohort study. BMC Cancer 2014;14:728.

6. Calle EE, Rodriguez C, Walker-Thurmond K, et al. Overweight, obesity, and mortality from cancer in a prospectively studied cohort of U.S. adults. N Engl J Med 2003;348:1625–38.

7. Eibl G, Cruz-Monserrate Z, Korc M, et al. Diabetes Mellitus and Obesity as Risk Factors for Pancreatic Cancer. J Acad Nutr Diet 2018;118:555–567.

8. Bracci PM. Obesity and pancreatic cancer: overview of epidemiologic evidence and biologic mechanisms. Mol Carcinog 2012;51:53–63.

9. Ng M, Fleming T, Robinson M, et al. Global, regional, and national prevalence of overweight and obesity in children and adults during 1980-2013: a systematic analysis for the Global Burden of Disease Study 2013. Lancet 2014;384:766–81.

10. Rossi EL, de Angel RE, Bowers LW, et al. Obesity-Associated Alterations in Inflammation, Epigenetics, and Mammary Tumor Growth Persist in Formerly Obese Mice. Cancer Prev Res (Phila) 2016;9:339–48.

11. Lashinger LM, Ford NA, Hursting SD. Interacting inflammatory and growth factor signals underlie the obesity-cancer link. J Nutr 2014;144:109–13.

12. Gomez-Chou S, Swidnicka-Siergiejko A, Badi N, et al. Lipocalin-2 Promotes Pancreatic Ductal Adenocarcinoma by Regulating Inflammation in the Tumor Microenvironment. Cancer Res 2017.

13. Wueest S, Konrad D. The controversial role of IL-6 in adipose tissue on obesity-induced dysregulation of glucose metabolism. Am J Physiol Endocrinol Metab 2020;319:E607–E613.

14. Klein AP. Pancreatic cancer epidemiology: understanding the role of lifestyle and inherited risk factors. Nat Rev Gastroenterol Hepatol 2021;18:493–502.

15. Keihanian T, Barkin JA, Souto EO. Early Detection of Pancreatic Cancer: Risk Factors and the Current State of Screening Modalities. Gastroenterol Hepatol (N Y) 2021;17:254–262.

16. Xu M, Jung X, Hines OJ, et al. Obesity and Pancreatic Cancer: Overview of Epidemiology and Potential Prevention by Weight Loss. Pancreas 2018;47:158–162.

17. Wadden TA, Tronieri JS, Butryn ML. Lifestyle modification approaches for the treatment of obesity in adults. Am Psychol 2020;75:235–251.

18. Kruk J, Czerniak U. Physical activity and its relation to cancer risk: updating the evidence. Asian Pac J Cancer Prev 2013;14:3993–4003.

19. Matthews CE, Moore SC, Arem H, et al. Amount and Intensity of Leisure-Time Physical Activity and Lower Cancer Risk. J Clin Oncol 2020;38:686–697.

20. Na HK, Oliynyk S. Effects of physical activity on cancer prevention. Ann N Y Acad Sci 2011;1229:176–83.

21. Hsueh HY, Pita-Grisanti V, Gumpper-Fedus K, et al. A review of physical activity in pancreatic ductal adenocarcinoma: Epidemiology, intervention, animal models, and clinical trials. Pancreatology 2022;22:98–111.

22. Wang Q, Zhou W. Roles and molecular mechanisms of physical exercise in cancer prevention and treatment. J Sport Health Sci 2021;10:201–210.

23. Gonzalez-Gil AM, Elizondo-Montemayor L. The Role of Exercise in the Interplay between Myokines, Hepatokines, Osteokines, Adipokines, and Modulation of Inflammation for Energy Substrate Redistribution and Fat Mass Loss: A Review. Nutrients 2020;12.

24. Farris MS, Mosli MH, McFadden AA, et al. The Association between Leisure Time Physical Activity and Pancreatic Cancer Risk in Adults: A Systematic Review and Meta-analysis. Cancer Epidemiol Biomarkers Prev 2015;24:1462–73.

25. Philip B, Roland CL, Daniluk J, et al. A high-fat diet activates oncogenic Kras and COX2 to induce development of pancreatic ductal adenocarcinoma in mice. Gastroenterology 2013;145:1449–58.

26. Percie du Sert N, Hurst V, Ahluwalia A, et al. The ARRIVE guidelines 2.0: updated guidelines for reporting animal research. BMJ Open Sci 2020;4:e100115.

27. Jackson EL, Willis N, Mercer K, et al. Analysis of lung tumor initiation and progression using conditional expression of oncogenic K-ras. Genes Dev 2001;15:3243–8.

28. Ji B, Song J, Tsou L, et al. Robust acinar cell transgene expression of CreErT via BAC recombineering. Genesis 2008;46:390–5.

29. Olive KP, Jacobetz MA, Davidson CJ, et al. Inhibition of Hedgehog signaling enhances delivery of chemotherapy in a mouse model of pancreatic cancer. Science 2009;324:1457–61.

30. Ma Y, Hwang RF, Logsdon CD, et al. Dynamic mast cell-stromal cell interactions promote growth of pancreatic cancer. Cancer Res 2013;73:3927–37.

31. Huang W, Liu X, Queen NJ, et al. Targeting Visceral Fat by Intraperitoneal Delivery of Novel AAV Serotype Vector Restricting Off-Target Transduction in Liver. Mol Ther Methods Clin Dev 2017;6:68–78.

32. Liu X, Magee D, Wang C, et al. Adipose tissue insulin receptor knockdown via a new primate-derived hybrid recombinant AAV serotype. Mol Ther Methods Clin Dev 2014;1:8-.

33. Bergin SM, Xiao R, Huang W, et al. Environmental activation of a hypothalamic BDNF-adipocyte IL-15 axis regulates adipose-natural killer cells. Brain Behav Immun 2021;95:477–488.

34. Xiao R, Bergin SM, Huang W, et al. Enriched environment regulates thymocyte development and alleviates experimental autoimmune encephalomyelitis in mice. Brain Behav Immun 2019;75:137–148.

35. Hruban RH, Adsay NV, Albores-Saavedra J, et al. Pathology of genetically engineered mouse models of pancreatic exocrine cancer: consensus report and recommendations. Cancer Res 2006;66:95–106.

36. Berman-Booty LD, Sargeant AM, Rosol TJ, et al. A review of the existing grading schemes and a proposal for a modified grading scheme for prostatic lesions in TRAMP mice. Toxicol Pathol 2012;40:5–17.

37. Raudvere U, Kolberg L, Kuzmin I, et al. g:Profiler: a web server for functional enrichment analysis and conversions of gene lists (2019 update). Nucleic Acids Res 2019;47:W191–W198.

38. Lauby-Secretan B, Scoccianti C, Loomis D, et al. Body Fatness and Cancer--Viewpoint of the IARC Working Group. N Engl J Med 2016;375:794–8.

39. Park SK, Jung JY, Oh CM, et al. Daily Vigorous Intensity Physical Activity and Its Preventive Effect on Pancreatic Cancer. Cancer Res Treat 2021.

40. Schmid D, Leitzmann MF. Association between physical activity and mortality among breast cancer and colorectal cancer survivors: a systematic review and meta-analysis. Ann Oncol 2014;25:1293–311.

41. Florez Bedoya CA, Cardoso ACF, Parker N, et al. Exercise during preoperative therapy increases tumor vascularity in pancreatic tumor patients. Sci Rep 2019;9:13966.

42. Gupta P, Hodgman CF, Alvarez-Florez C, et al. Comparison of three exercise interventions with and without gemcitabine treatment on pancreatic tumor growth in mice: No impact on tumor infiltrating lymphocytes. Front Physiol 2022;13:1039988.

43. Katsourakis A, Vrabas I, Dimitriadis C, et al. How Exercise Can Influence Oxidative Stress and Glucose Levels after Pancreatic Resection: A Randomised Controlled Trial. Dig Surg 2020;37:205–210.

44. Pannala R, Leirness JB, Bamlet WR, et al. Prevalence and clinical profile of pancreatic cancer-associated diabetes mellitus. Gastroenterology 2008;134:981–7.

45. Sharma A, Smyrk TC, Levy MJ, et al. Fasting Blood Glucose Levels Provide Estimate of Duration and Progression of Pancreatic Cancer Before Diagnosis. Gastroenterology 2018;155:490–500 e2.

46. Ciapaite J, van den Broek NM, Te Brinke H, et al. Differential effects of short- and long-term high-fat diet feeding on hepatic fatty acid metabolism in rats. Biochim Biophys Acta 2011;1811:441–51.

47. Pranprawit A, Wolber FM, Heyes JA, et al. Short-term and long-term effects of excessive consumption of saturated fats and/or sucrose on metabolic variables in Sprague Dawley rats: a pilot study. J Sci Food Agric 2013;93:3191–7.

48. Bowers LW, Doerstling SS, Shamsunder MG, et al. Reversing the Genomic, Epigenetic, and Triple-Negative Breast Cancer-Enhancing Effects of Obesity. Cancer Prev Res (Phila) 2022;15:581–594.

49. Dasso NA. How is exercise different from physical activity? A concept analysis. Nurs Forum 2019;54:45–52.

50. Gomes-Santos IL, Amoozgar Z, Kumar AS, et al. Exercise Training Improves Tumor Control by Increasing CD8(+) T-cell Infiltration via CXCR3 Signaling and Sensitizes Breast Cancer to Immune Checkpoint Blockade. Cancer Immunol Res 2021;9:765–778.

51. Wennerberg E, Lhuillier C, Rybstein MD, et al. Exercise reduces immune suppression and breast cancer progression in a preclinical model. Oncotarget 2020; 11:452–461.

52. Farhood B, Najafi M, Mortezaee K. CD8(+) cytotoxic T lymphocytes in cancer immunotherapy: A review. J Cell Physiol 2019;234:8509–8521.

53. Masugi Y, Abe T, Ueno A, et al. Characterization of spatial distribution of tumor-infiltrating CD8(+) T cells refines their prognostic utility for pancreatic cancer survival. Mod Pathol 2019;32:1495–1507.

54. Hou YC, Chao YJ, Hsieh MH, et al. Low CD8(+) T Cell Infiltration and High PD-L1 Expression Are Associated with Level of CD44(+)/CD133(+) Cancer Stem Cells and Predict an Unfavorable Prognosis in Pancreatic Cancer. Cancers (Basel) 2019;11.

55. Kurz E, Hirsch CA, Dalton T, et al. Exercise-induced engagement of the IL-15/IL-15Ralpha axis promotes anti-tumor immunity in pancreatic cancer. Cancer Cell 2022.

56. Guo S, Huang Y, Zhang Y, et al. Impacts of exercise interventions on different diseases and organ functions in mice. J Sport Health Sci 2020;9:53–73.

57. Park J, Euhus DM, Scherer PE. Paracrine and endocrine effects of adipose tissue on cancer development and progression. Endocr Rev 2011;32:550–70.

58. Saha A, Ahn S, Blando J, et al. Proinflammatory CXCL12-CXCR4/CXCR7 Signaling Axis Drives Myc-Induced Prostate Cancer in Obese Mice. Cancer Res 2017;77:5158–5168.

59. Okumura T, Ohuchida K, Sada M, et al. Extra-pancreatic invasion induces lipolytic and fibrotic changes in the adipose microenvironment, with released fatty acids enhancing the invasiveness of pancreatic cancer cells. Oncotarget 2017;8:18280–18295.

60. Dirat B, Bochet L, Dabek M, et al. Cancer-associated adipocytes exhibit an activated phenotype and contribute to breast cancer invasion. Cancer Res 2011;71:2455–65.

