## Supplementary Materials and Methods for "Physical Activity Delays Obesity-Associated Pancreatic Ductal Adenocarcinoma in Mice and Decreases Inflammation"

*Immunohistochemistry*

Paraffin tissue sections (4 um) were deparaffinized and treated with Cytomation Target Retrieval solution (pH 6.0, Dako, Glostrup, Denmark) in a Decloaking Chamber (Biocare Medical, Concord, CA) heated to 125 °C, then cooled to 90 °C for 10 seconds before cooling with the lid removed for 10 minutes. Slides were transferred to a Dako Universal Training Center automatic immunostainer. After hydrogen peroxide treatment and serum-free blocking, slides were stained for macrophages, proliferation, smooth muscle actin, with their respective secondary antibodies. (**Supplementary Table 3**) All slides were incubated with Vector RTU ABC elite complex (Vector Laboratores) followed by diaminobenzidine chromagen then counterstained with hematoxylin. Negative controls included isotype-matched controls using nonspecific IgG at similar concentrations. Images were captured from whole-slide images acquired with an Aperio XT slide scanner (Leica Biosystems, Buffalo Grove, IL) using a 20X objective.

*ELISA and Luminex Multiplex Cytokine Analysis*

Human and mouse LCN2 Quantikine ELISA kits (R&D Systems, Minneapolis, MN) were used to measure serum LCN2 levels following manufacturer's recommendations. A panel of 32 cytokines, chemokines, and growth factors was measured in mice serum using a Luminex Multiplex Cytokine Kit (Procarta Cytokine Assay, eBioscience, San Diego, CA). A panel of 21 cytokines, chemokines, and growth factors was measured in human serum using a MilliPlex Multiplex Assay (Millipore Sigma, Burlington, MA). Samples were analyzed in duplicate and quantified using analyte-specific standard curves for each batch.

*RNA isolation and quantitative RT-PCR*

Mouse adipose tissue RNA was isolated using TRIzol reagent (Life Technologies). cDNA was generated using the Verso cDNA synthesis kit (ThermoFisher Scientific, Waltham, MA). Quantitative RT-PCR (ThermoFisher Scientific) was performed using mouse primers for IL-15 (forward 5'-GTAGGTCTCCCTAAAACAGAGGC-3', reverse 5'-TCCAGGAGAAAGCAGTTCATTGC-3'), and IL-15 receptor alpha (ra) (forward 5-'CGTGTCCACCTCCCGTATCTA-3', reverse 5'-AGACATACCTCTCCCTGGAGT-3') and normalized to GAPDH (forward 5'-AGCCTCGTCCCGTAGACAAAA-3', reverse 5'-GCC TTGACTGTGCCGTTGATT-3') (Integrated DNA Technologies IDT, Coralville, IA).

*Affymetrix gene analysis*

Mouse adipose tissue RNA was cleaned using the RNeasy MinElute Cleanup Kit (Qiagen, Hilden, Germany) followed by reprecipitation prior to Mouse Clariom S Affymetrix analysis. Data was analyzed using the Transcriptome Analysis Console after log transformation and Gene Ontology analysis with the g:Profiler platform[1].

Supplementary references

1. Raudvere U, Kolberg L, Kuzmin I, et al. g:Profiler: a web server for functional enrichment analysis and conversions of gene lists (2019 update). Nucleic Acids Res 2019;47:W191-W198.

| Lesion Grade | Diagnosis | Distribution | Adjusted Lesion Score |
|---|---|---|---|
| 0 | Normal | Diffuse | 0 |
| 1 | mPanIN-1 | Focal (F) | 1 |
| 1 | mPanIN-1 | Multifocal (MF) | 2 |
| 1 | mPanIN-1 | Diffuse | 3 |
| 2 | mPanIN-2 | Focal (F) | 4 |
| 2 | mPanIN-2 | Multifocal (MF) | 5 |
| 2 | mPanIN-2 | Diffuse | 6 |
| 3 | mPanIN-3 | Focal (F) | 7 |
| 3 | mPanIN-3 | Multifocal (MF) | 8 |
| 3 | mPanIN-3 | Diffuse | 9 |
| 4 | PDAC | Focal (F) | 10 |
| 4 | PDAC | Multifocal (MF) | 11 |
| 4 | PDAC | Diffuse | 12 |

**Supplementary Table 1. Summary of Mouse Pancreas Adjusted Lesion Scoring System.** This grading scheme incorporates both the most severe and most common lesions, both of which contribute to the pathology present within a section. The grades range from 0-4 with normal pancreas as the least severe (grade 0) and pancreatic ductal carcinoma the most severe lesion (grade 4). The first grade accounts for the most severe lesion within the section. The second grade, using the same scale, accounts for the most common lesion in the section. After the most severe and the most common lesions within a section are identified, the distributions of these lesions are determined and described as focal if less than 3 foci contain the lesion, multifocal if there are three or more foci or less than 50% of the section containing the lesion or diffuse if greater than 50% of section contains the lesion. Adjusting the lesion grades to include an indication of distribution provides two adjusted scores, one for most severe lesion, and the other for most common ranging from 0 (normal) to 12 (diffuse PDA). These adjusted scores are then added to obtain a sum that reflects the most severe lesion and its distribution and the most common lesion and its distribution (sum of the adjusted lesion scores) which were then added with the fibrosis and inflammation score to generate the total pathology score.

| Fibrosis Score | | Inflammation score | |
|---|---|---|---|
| 0 | Normal and no fibrosis | 0 | No inflammation |
| 1 | 1-10% increase in collagenous stroma and occasional; periductular or scant perilobular fibrosis | 1 | Minimal infiltration of periductal tissue and occasional scattered leukocytes |
| 2 | 10-30% increase in collagenous stroma showing moderate fibrosis | 2 | Inflammation around ducts and extending into the parenchyma (< 50% of lobules) with moderate numbers of leukocytes |
| 3 | 30-50% increase in collagenous stroma showing extensive fibrosis | 3 | Inflammation around ducts and extending into the parenchyma (51%-75% of lobules) with dense aggregates of leukocytes |
| 4 | More than 50% fibrosis | 4 | Inflammation around ducts and extending into the parenchyma (>75% lobules) |

**Supplementary Table 2. Summary of Fibrosis and Inflammation Scoring System.**

| Antibody | Antibody source | Antibody dilution | Secondary antibody information | Application |
|---|---|---|---|---|
| 647 F4/80 | Clone BM8; (Biolegend, San Diego, CA) | 1:50 | NA | Flow cytometry |
| FITC CD11c | Clone B-ly6 (Biolegend) | 1:50 | NA | Flow cytometry |
| PE/Cy7 CD11b | Clone M1/70 (Biolegend) | 1:50 | NA | Flow cytometry |
| PE Ly-6G | Clone YTS156.7.7 (Biolegend) | 1:50 | NA | Flow cytometry |
| 488 Ly6C | Clone HK1.4 (Biolegend) | 1:50 | NA | Flow cytometry |
| 647 CD3 | Clone 17A2 (Biolegend) | 1:50 | NA | Flow cytometry |
| PE NK1.1 | Clone S17016D (Biolegend) | 1:50 | NA | Flow cytometry |
| 488 CD4 | Clone RM4-5 (Biolegend) | 1:50 | NA | Flow cytometry |
| PE/Cy7 CD8 | Clone 53-6.7 (Biolegend) | 1:50 | NA | Flow cytometry |
| F4/80 | MCA497G rat monoclonal (Serotec) | 1:100 | Biotinylated rabbit anti-rat (Vector Laboratories) 1:200 dilution. | Immunohistochemistry |
| Ki67 | RM-9106-R7 rabbit monoclonal (Thermo Scientific) | 1:100 | Biotinylated goat anti-rabbit (Vector Laboratories) 1:200 dilution. | Immunohistochemistry |
| αSMA | M0851 mouse monoclonal [clone 1A4] (Dako) | 1:50 | Mouse-on-Mouse Polymer (Biocare Medical) | Immunohistochemistry |

**Supplementary Table 3. Summary of Antibodies Utilized.**

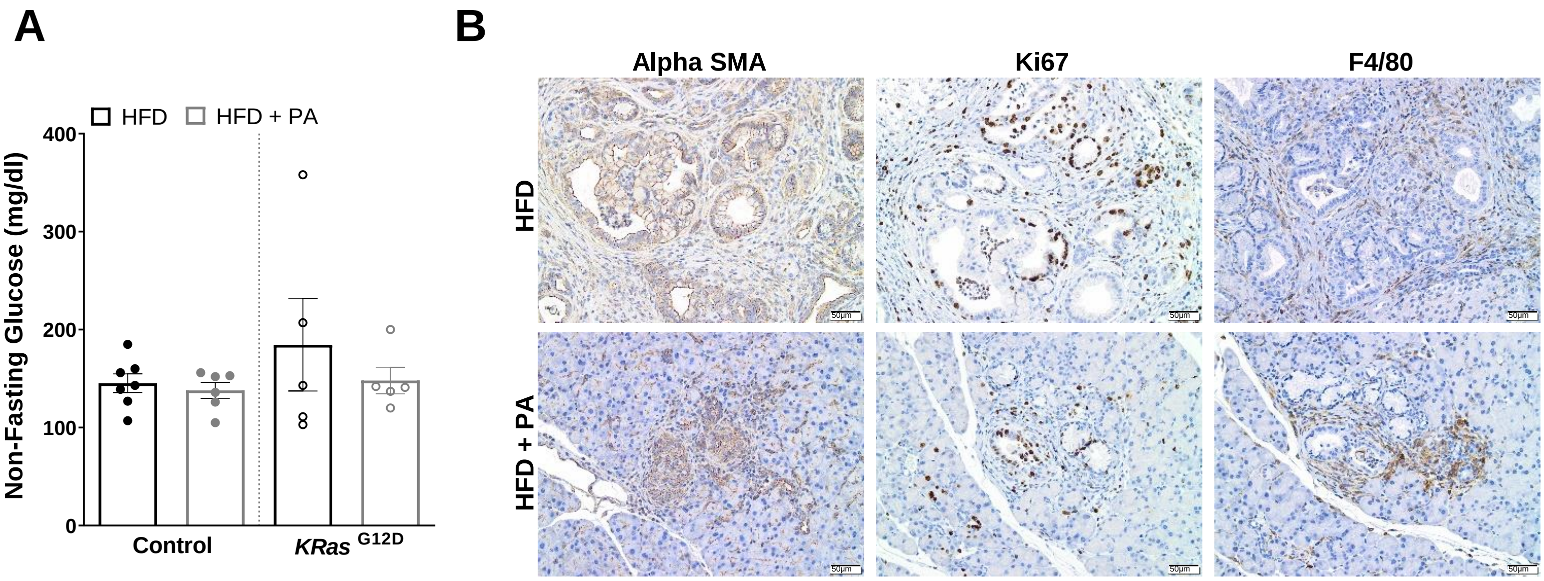


**Supplementary Figure 1. PA does not modulate non-fasting glucose, but delays fibrosis, proliferation, and macrophage infiltration in an obesity-induced GEMM of PDAC. (A)** Non-fasting blood glucose after 31 days of HFD or HFD + PA intervention described in Figure 2A, analyzed with two-way ANOVA. **(B)** Representative IHC stains of fibrosis (αSMA), proliferation (Ki67) and macrophages (F4/80) in the pancreas of Kras$^{G12D}$ mice described in Figure 2A (original magnification=20x, scale bar=50μm).

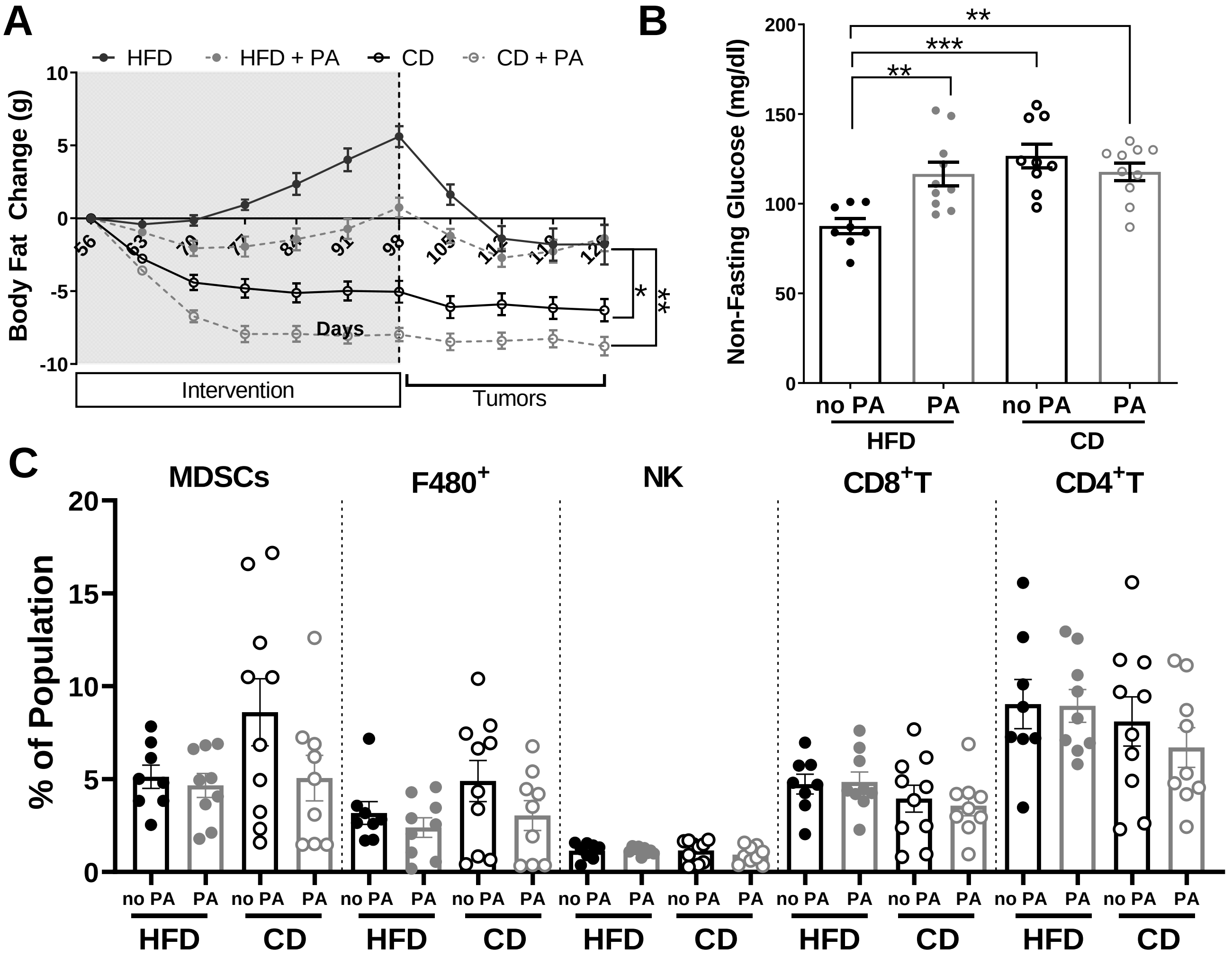


**Supplementary Figure 2. CD and PA intervention reduce body weight, increase non-fasting glucose levels, and do not modulate immune cell populations in an orthotopic mouse model of PDAC.** (**A**) Weekly change in body fat over time, analyzed with mixed-effects analysis, comparing to HFD control (**B**) Average non-fasting glucose levels at end-point, analyzed via two-way ANOVA. **(C)** Flow cytometry analysis of splenic MDSCs, F480$^{+}$ Macrophages, NK cells, CD4$^{+}$ T cells, and CD8$^{+}$ T cells, analyzed via two-way ANOVA test.

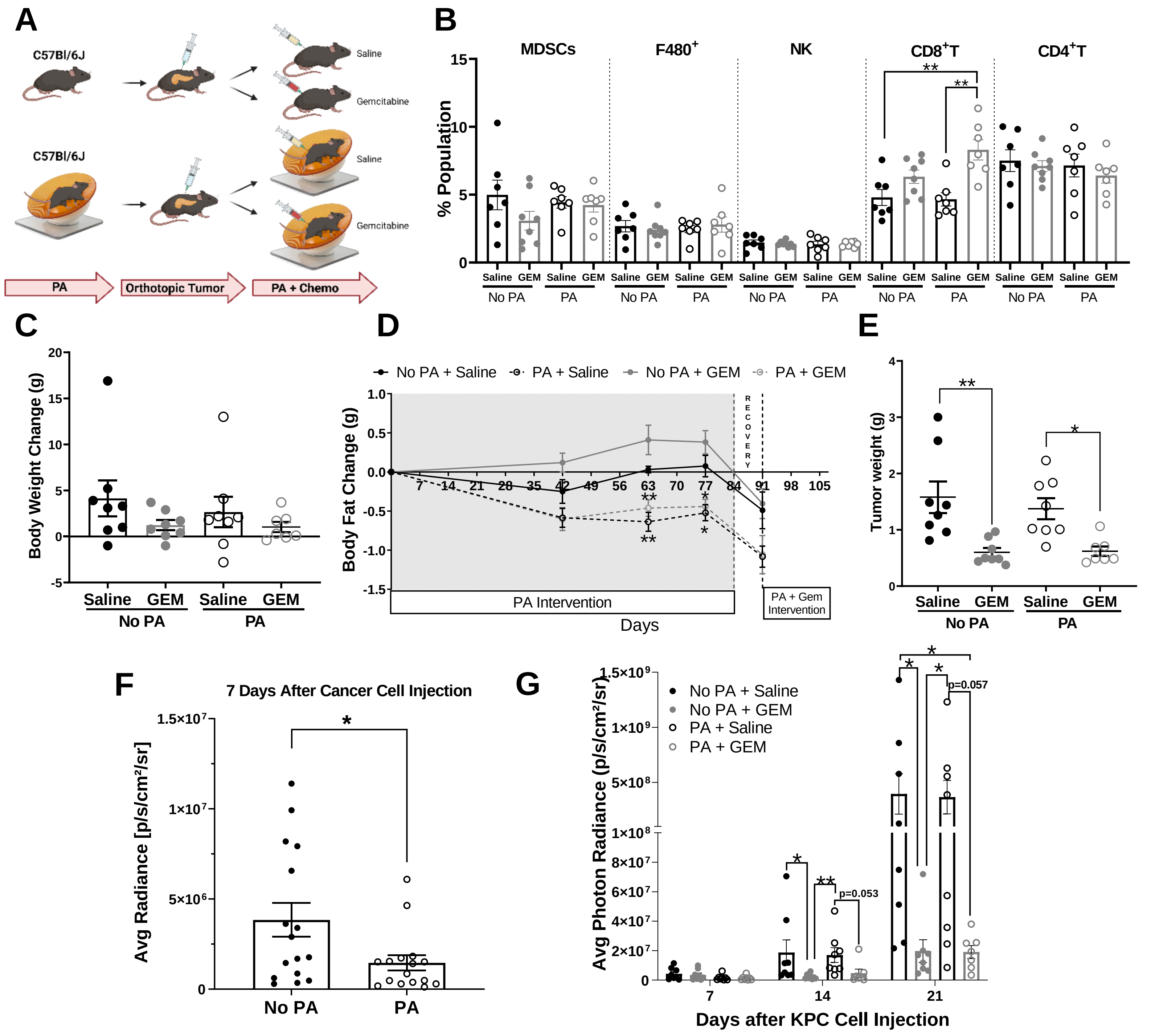


**Supplementary Figure 3. PA delays tumor growth only 7 days after cancer cell injections, but this effect is not sustained throughout time, when combine with chemotherapy, PA increases CD8$^{+}$T cells**. (**A**) At 104 days of age, male C57BL/6J mice were randomly assigned to either a PA or no PA intervention for 83 days before receiving orthotopic injections of cancer cells. Seven days after the orthotopic injections the mice received either saline or 33 mg/kg of gemcitabine chemotherapy twice a week (n= 8 per group). (**B**) Percent population of MDSCs, F480+ Macrophages, NK cells, CD8+ T cells, and CD4+ T cells, analyzed via two-way ANOVA. (**C**) Body weight change from day 0 to the end of the intervention, analyzed via two-way ANOVA compared to No PA + Saline. (**D**) Body fat change over time analyzed via two-way ANOVA. (**E**) Tumor weight at the end of the intervention, analyzed via two-way ANOVA after log-transformation. (**F**) Average tumor radiance seven days after orthotopic injections, analyzed after log transformation via unpaired t-test. (**G**) Average photon radiance of the pancreas per week, per group after orthotopic injections, analyzed via mixed-effects analysis after log transformation.
